# Quantifying the relationship between cell proliferation and morphology during development of the face

**DOI:** 10.1101/2023.05.12.540515

**Authors:** Rebecca M. Green, Lucas D. Lo Vercio, Andreas Dauter, Elizabeth C. Barretto, Jay Devine, Marta Vidal-García, Marta Marchini, Samuel Robertson, Xiang Zhao, Anandita Mahika, M. Bilal Shakir, Sienna Guo, Julia C. Boughner, Wendy Dean, Arthur D. Lander, Ralph S. Marcucio, Nils D. Forkert, Benedikt Hallgrímsson

## Abstract

Morphogenesis requires highly coordinated, complex interactions between cellular processes: proliferation, migration, and apoptosis, along with physical tissue interactions. How these cellular and tissue dynamics drive morphogenesis remains elusive. Three dimensional (3D) microscopic imaging poses great promise, and generates elegant images. However, generating even moderate through-put quantified images is challenging for many reasons. As a result, the association between morphogenesis and cellular processes in 3D developing tissues has not been fully explored. To address this critical gap, we have developed an imaging and image analysis pipeline to enable 3D quantification of cellular dynamics along with 3D morphology for the same individual embryo. Specifically, we focus on how 3D distribution of proliferation relates to morphogenesis during mouse facial development. Our method involves imaging with light-sheet microscopy, automated segmentation of cells and tissues using machine learning-based tools, and quantification of external morphology via geometric morphometrics. Applying this framework, we show that changes in proliferation are tightly correlated to changes in morphology over the course of facial morphogenesis. These analyses illustrate the potential of this pipeline to investigate mechanistic relationships between cellular dynamics and morphogenesis during embryonic development.

## INTRODUCTION

Disentangling the mechanisms of morphogenesis that translate the behaviour of individual cells to an intricate three-dimensional (3D) organismal form has been a major objective of embryology and developmental biology since the early 19^th^ century. The cellular basis for morphogenesis is complex, involving spatio-temporal variation in cell proliferation, apoptosis, adhesion, and polarity. Morphogenesis also involves interactions between cells, the extracellular matrix, and mechanical forces that emanate from surrounding tissues and the extra-embryonic environment (Boehm et al., 2010; Seilacher, 1991; Newman and Comper, 1990; Niessen et al., 2011; Davies, 2013). Connecting these mechanisms to 3D change in organismal form requires quantification of cellular processes in whole embryonic structures over developmental time. While three dimensional morphology can be quantified from microCT and optical projection tomography images (Parsons et al., 2008; Boughner et al., 2008; Xu et al., 2015; Martínez-Abadías et al., 2018), cellular dynamics are much less accessible in three dimensions. This is because the vast majority of quantification of cellular dynamics is based on analysis of serial histological sections, with localized sampling rather than whole embryonic structures (Ramaesh and Bard, 2003; Miklius and Hilgenfeldt, 2011; Russ and Dehoff, 2012). Serial histological sections are rife with artifacts ranging from distortions from fixation and sectioning (Xiao et al., 2010)) to plane of section artifacts that occur when complex 3D structures are reduced to two dimensions (Russ and Dehoff, 2012). Serial sectioning of whole embryonic structures is also labour- and time-intensive, which makes this method unsuitable for analysis of large samples. However, recent advances in 3D imaging of whole tissue samples, including entire embryos, as well the ability to combine such images with multiple molecular markers is now creating the opportunity for true quantitative integration of cellular dynamics and morphology. These imaging modalities also generate large and complex datasets that demand novel image processing and analysis methods. In this study, we deploy a novel imaging and image analysis pipeline to examine cell proliferation in the developing face, thereby advancing the capacity for quantitative integration of cellular dynamics and morphology.

The development of the vertebrate face involves directional outgrowth and fusion of distinct facial prominences that form different facial regions. In mammals the upper jaw forms from divisions of the frontonasal prominence along the midline - the medial and lateral nasal prominences - fusing together and with the maxillary prominence (Chai and Maxson Jr, 2006). The facial prominences must grow in a coordinated manner that includes both outgrowth and alignment to allow for fusion (Chai and Maxson Jr, 2006; Green et al., 2015). While the processes involved in outgrowth of structures such as limb buds have been fairly well characterized (Boehm et al., 2010), these studies have been more difficult to perform in the face due to the complexity of signaling pathway interactions and morphology. Unlike the limb, outgrowth of the facial prominences is orchestrated by multiple signaling centers, including around the frontonasal prominence (WNT, FGFs), in the leading edge of the maxilla (FGFs, BMPs), and in the forebrain (SHH, BMPs) (Marchini et al., 2021). These signals drive a combination of local cell proliferation, migration of neural crest cells, and additionally likely affect mechanical properties of the tissue. The complex interplay of growth, cell migration, and signaling has made it difficult to tease apart the roles of individual factors. While there is a general intuition that spatiotemporal regulation of cell proliferation within the facial prominences is important in face formation, this regulation has not been shown quantitatively. Further, the role of proliferation in relation to other potential processes, such as apoptosis, adhesion, polarity, or mechanical influences from the epithelium, needs to be understood in much finer detail to explain facial development and account for congenital facial malformations, such as orofacial clefts.

Recent advances in tissue clearing methods coupled with light-sheet fluorescence microscopy (LSFM) allow visualization of individual cells within whole embryos or anatomical structures (Weber et al., 2014; Yue et al., 2020; Udan et al., 2014; Elisa et al., 2018). This visualization enables quantification of cellular level variation as well as the morphology of the developing tissues at the level of individual embryos (McDole et al., 2018; Hobson et al., 2022). Accordingly, cellular dynamics such as proliferation, orientation, apoptosis, cell size, and cell density can be related quantitatively to variation in morphology. Importantly, these analyses can be performed on individuals, as opposed to group-wise, allowing for cellular-level investigations of the mechanisms underlying among-individual variation in morphology. This individual-level investigation is critical for elucidating mechanisms for genetically complex malformations or those which have variable penetrance or expressivity (Hallgrimsson et al., 2019). However, even with light-sheet microscopy, this task is challenging. In mice, for example, tissue density precludes single-cell visualization without clearing to allow light penetration after embryonic day (E) 8.5 (McDole et al., 2018). Quantifying variation among embryos requires some form of image registration to identify homology between corresponding anatomical locations. However, the large amount of shape changes characteristic of morphogenesis complicates the construction of atlas-based registration pipelines without copious quantities of data (Wong et al., 2015). LSFM images are prone to artifacts from optical aberrations or deviations in refractive index across a specimen. These issues present challenges for developing segmentation protocols, either human or computational, as a region of an image may be out of focus. Further, the images files are large, ranging from 300 MB to 1 TB, depending on embryo size and image resolution. Therefore, LSFM images require extensive computational resources for data management and analysis. Overcoming these obstacles has been challenging with available tools and conventional methods.

As such, previous work towards this goal has been hampered by both limitations in imaging technology as well as image processing and informatics. Optical projection tomography (OPT) has been used to create anatomical images and quantitative analyses for gene expression (Sharpe et al., 2002) as well as markers of cellular dynamics (Boehm et al., 2010). The limitation here is that, as the anatomical structures to be imaged get larger, the effective resolution of OPT imaging decreases, as this is ultimately dependent on the geometry and detection aperture (Wong et al., 2013; Liu et al., 2019). While early zebrafish embryos are sufficiently small to allow such imaging (Lindsey and Kaslin, 2017), structures such as whole mouse embryo heads are too large to allow cellular level imaging with currently available OPT systems at the larger ages. Light-sheet microscopy overcomes this limitation of OPT using tiling and laser optics by illuminating a thin sheet of the sample (Olarte et al., 2018). This method also reduces photodamage, which can occur in confocal fluorescence microscopy (Olarte et al., 2018). The image processing and imaging informatics gap is that all of these methods generate large and noisy volumetric image sets that require extensive post-processing and advanced registration methods. Our work addresses this gap using advanced image processing methods.

In a previous paper, we (Lo Vercio et al., 2022) demonstrated that Convolutional Neural Networks (CNNs), particularly U-net architectures, can efficiently segment the mesenchyme (Mes) and neural ectoderm (NE) in nuclei-stained 2D LSFM images. This type of deep learning architecture has also been used for other aspects of light-sheet data processing (Yin et al., 2022; Hallou et al., 2021). Furthermore, CNNs efficiently segmented cells labeled for a proliferation marker in LSFM images of E9.5-10.5 embryos. Here, we leverage this advanced image processing approach to develop and apply a method to quantify the relationship between cellular dynamics and 3D morphology in embryonic tissues based on light sheet microscopy. We first apply our previously developed tools for segmenting tissues and cells at the individual level (Lo Vercio et al., 2022). Next, a 3D image registration pipeline technique (Devine et al., 2020) is employed to create 3D atlases that separate neural and non-neural tissues and contain volumetric representations of cell proliferation. We then apply the tools of geometric morphometrics to quantify spatiotemporal variation in cell proliferation along with embryonic surface morphology. Specifically, we seek to determine a) the relationship between proliferation and morphogenesis, and b) the contribution of anatomical variation in proliferation to among-individual variation in cranial morphology. Successful quantitative integration of cellular dynamics is critical for unravelling the mechanisms of morphogenesis, but also the mechanistic basis for among-individual variation, including the etiology of structural birth defects.

## RESULTS

### Light-sheet imaging and workflow to build atlases of shape, tissues and cellular dynamics from mouse embryos

LSFM imaging of mouse facial tissue can be used to generate both cellular level volumetric data and a quantifiable 3D exterior surface (Fig. 1). Building on previous work (Lo Vercio et al., 2022), we established an automatic workflow for segmenting, registering, and quantifying cellular dynamics from LSFM scans of mouse embryos (Fig. 2). Here, we scanned 20 wildtype C57BL/6J mouse embryos ranging in age from embryonic day (E) 10 to 11.5. This age range encompasses early facial morphogenesis and captures the cell dynamics involved in facial prominence outgrowth. The embryos were separated into half-day groups based on tail somite number, with five embryos per group. Half-day intervals were originally chosen to compose well-matched groups that can be used for atlas generation. We produced registered volumetric image sets from these scans and analyzed spatiotemporal patterning of proliferation in whole embryonic heads.

**Fig. 1.**
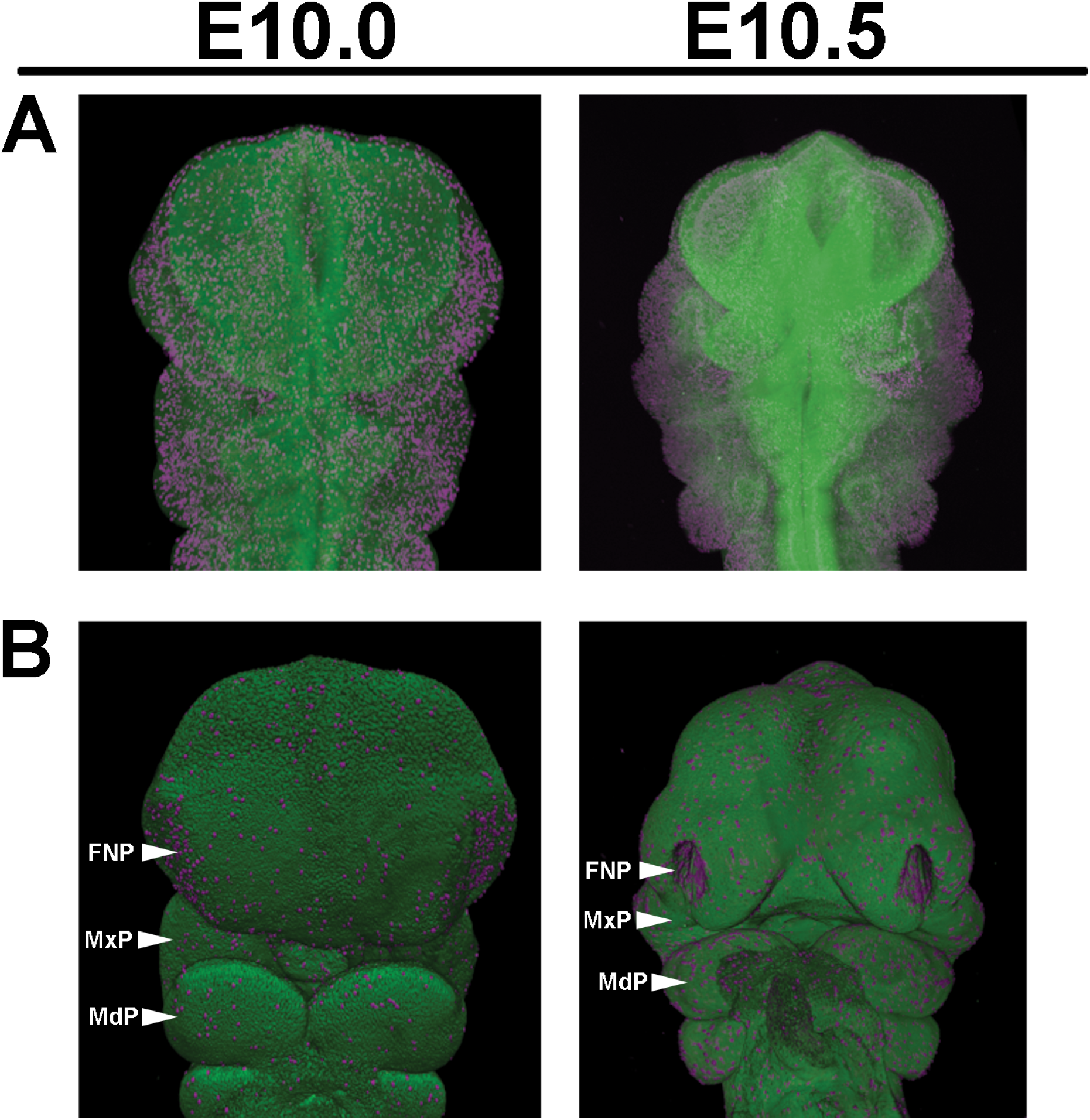
Maximum projection intensity images in which Nuclear Green has been used to stain nuclei; phospho-Histone H3 (pHH3) has been stained in magenta to mark cell proliferation. Embryos were harvested at E10-11 from C57Bl/6J mice following timed matings. Embryos were cleared and prepared following the CUBIC protocol, antibody stained, and mounted in 1.5% low-melt agarose. Images were captured on a Zeiss Lightsheet Z1 microscope and analyzed in Arivis. Embryo staging was established by counting tail somites. FNP = Frontonasal Prominence, MxP = Maxillary Prominence, MdP = Mandibular Prominence Representative images of *n* > 20 samples.

**Fig. 2.**
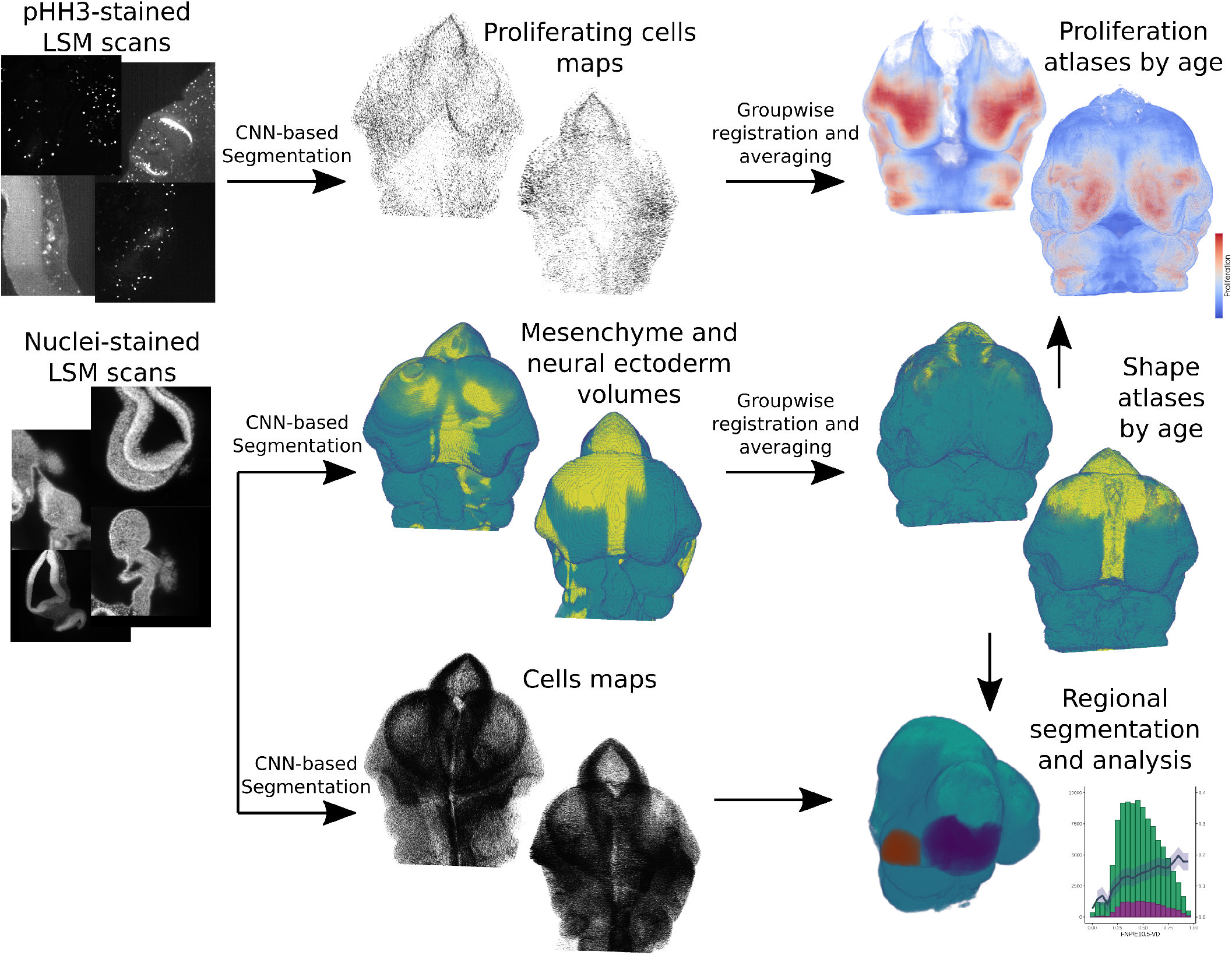
Overview of the workflow for creating shape and proliferation atlases from LSFM scans. Twenty mouse embryos are stained for nuclei (Nuclear Green) and proliferating cells (pHH3). For each specimen, proliferating cell maps and tissues are automatically segmented using CNNs (mesenchyme in teal, neural ectoderm in yellow). For each age group (E10, E10.5, E11, and E11.5), the groupwise affine-registration is performed using the tissue segmentation volume. Then, the resulting transformation for each sample is applied to its proliferating cell map. The shape atlas per age is obtained by majority voting in each voxel. The proliferation atlases are produced by smoothing and normalizing each proliferating cell map, and then averaging the resulting heat maps of the age group.

An LSFM instrument generates data as a stack of 2D slices. Each slice is generated as the laser sheet passes through the sample. The within-slice resolution is typically 6-8 times higher than the between-slice resolution. Here, our analysis focuses on using the stack data: the tissues, total cells, and proliferating cells are segmented in the 2D images, then the anisotropic z-stacks of segmented images are converted to isotropic volumes (Fig. 2), middle column). A groupwise registration strategy based on the tissue segmentation can be used to create atlases of shapes and cellular dynamics (Fig. 2, last column). This pipeline and accompanying documentation is available from https://github.com/lucaslovercio/LSMprocessing.

### Shape, tissue, and proliferation atlases between E10.0 and E11.5

An atlas of neural and non-neural tissues was generated for each age group. Images of these atlases are shown in Fig. 3A top row. Our analysis focused on the non-neural tissues, primarily mesenchyme, as most of the key events happening during facial morphogenesis happen in mesenchymal tissues. As we ran into difficulties segmenting cells in the highly dense neural tissue, we did not pursue further analysis in this tissue. From here, mean proliferation was calculated for each group of specimens (Fig. 3A, bottom row). In general, we observed high levels of proliferation in the frontonasal, mandibular, and maxillary prominences - areas where shape change is happening at these points in development.

**Fig. 3.**
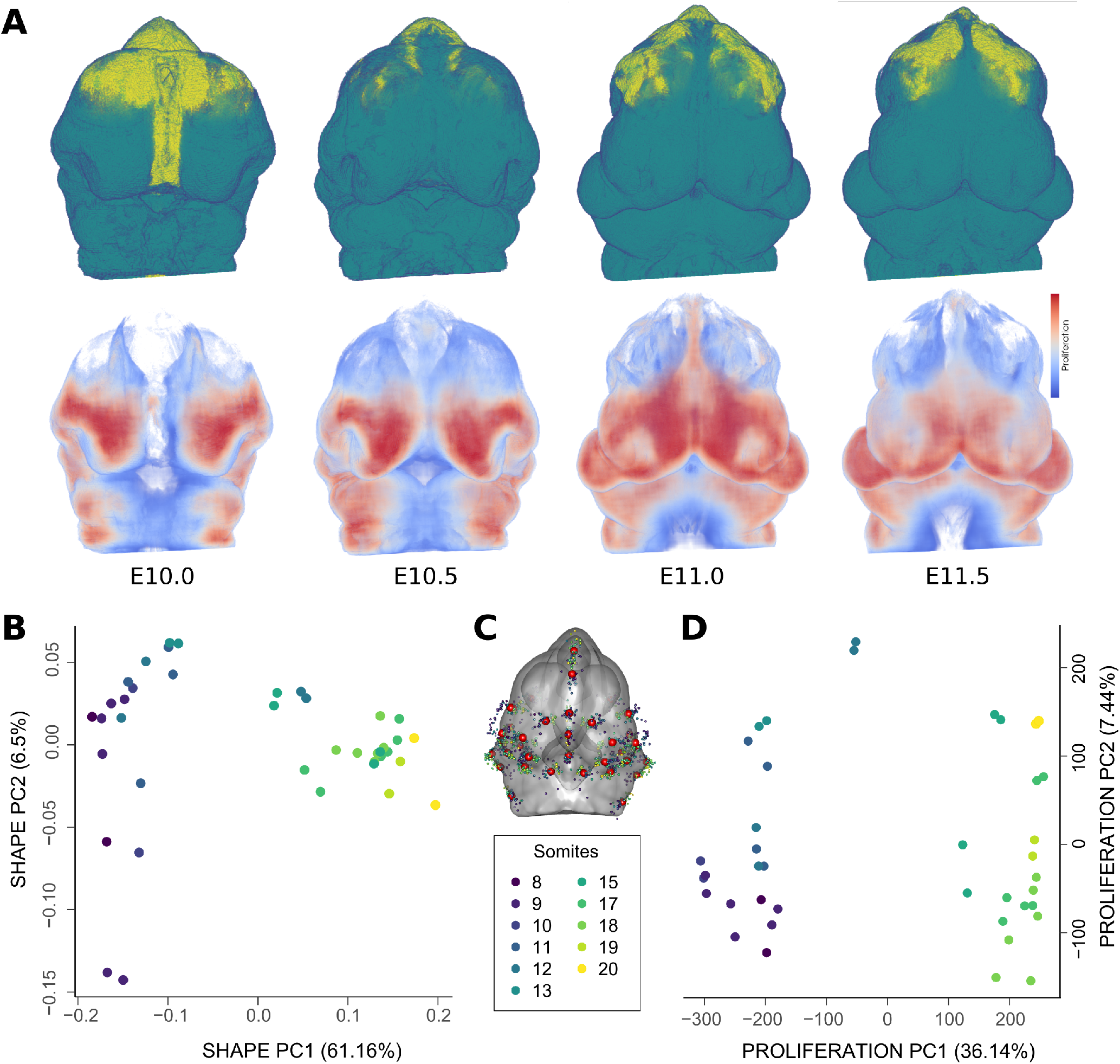
(A) Tissue and proliferation atlases for C57Bl/6J embryos between E10.0 and E11.5. The top row presents the tissue atlases for embryonic ages E10.0, E10.5, E11.0, and E11.5, showing mesenchyme in teal and neural ectoderm in yellow. Voxels are displayed slightly transparent for visualization purposes. The bottom row shows the proliferation map atlases for the same embryonic ages, only in the mesenchyme. (B,D) Principal Component Analysis (PCA) of Procrustes shape coordinates and facial proliferation, respectively, for the whole embryonic dataset. Specimens are coloured by tail somite count, as a proxy for age. Dataset comprises embryos from 8 to 20 somites. (C) Location of anatomical landmarks manually placed on the facial surfaces of embryonic samples in the atlas (average, in red). Dispersion of landmarks for each specimen is also shown and coloured by tail somite count.

In order to better understand the change in morphology and proliferation over the age series, we used landmark-based geometric morphometrics. We used 37 landmarks that are focused on the front of the face to capture the morphological shifts in the facial prominences (Fig. 3C) (Percival et al., 2014). To understand how the overall morphology of the embryo changes with time, we used Principal Component Analysis (PCA) to identify the largest axes of variance in our dataset. (Fig. 3B). For the PCA of facial shape coordinates, PC1 explained most of the variance (61.6%) and PC2 only 6.5% of the overall variance. Shape changes on PC1 were directly correlated to somite count, strongly separating samples with 8 to 14 somites (left region of the morphospace along PC1, negative values) from samples with 15 to 20 somites (right region of the morphospace along PC1, positive values). Interestingly, although we attempted to group the embryos using tail somite number, they still cluster morphologically into two distinct groups, E10-E10.5 and E11-E11.5. This clustering effect seems to be strongly driven by changes in the nasal prominence region. We found similar patterns on the PCA of proliferation in the front of the face, in which PC1 explained 36.14% of the overall variance, while PC2 only explained an additional 7.44% (Fig. 3D). We also found a strong separation of the same two age groups we observed in the shape PCA: samples with 8-14 somites (younger embryos) and samples with 15-20 somites (older embryos).

This observation raised the question if proliferation was uniform across these regions. We hypothesised that differences in proliferation density across developing facial prominences precede morphological change. To test this hypothesis, we manually segmented the frontonasal and maxillary prominences at the earlier time points (E10-E10.5) and calculated rates of proliferation across each developmental axis: anterior to posterior, dorsal to ventral, and distal to proximal. These tissues are highly proliferative and commonly perturbed in development, making them important morphological targets (Elmsie and Reardon, 1998). We calculated the absolute and proportional volume of proliferating nuclei compared to total nuclei across each axis of a prominence, averaging across all specimens in each group (Fig. 4). We found that the proportion of proliferating cells remains stable across most biological axes in these prominences. Additionally, there were no significant differences in mean proliferative fraction within each prominence, suggesting that the rate of proliferation remains similar in actively proliferating facial tissues. Tissue-specific means were similar, ranging between 14 and 18% of cells proliferating. We did observe a difference in the E10.5 frontonasal prominence along the dorsal-ventral axis, where cells proliferated more densely towards the dorsal side (interior) of the tissue (p < 0.0001). Notably, apparent patterns in proliferation may be related to the overall density of cells across tissues, which changed with time and prominence location. These apparent differences in cell densities could relate to morphological change over development and will need to be more carefully examined in future studies.

**Fig. 4.**
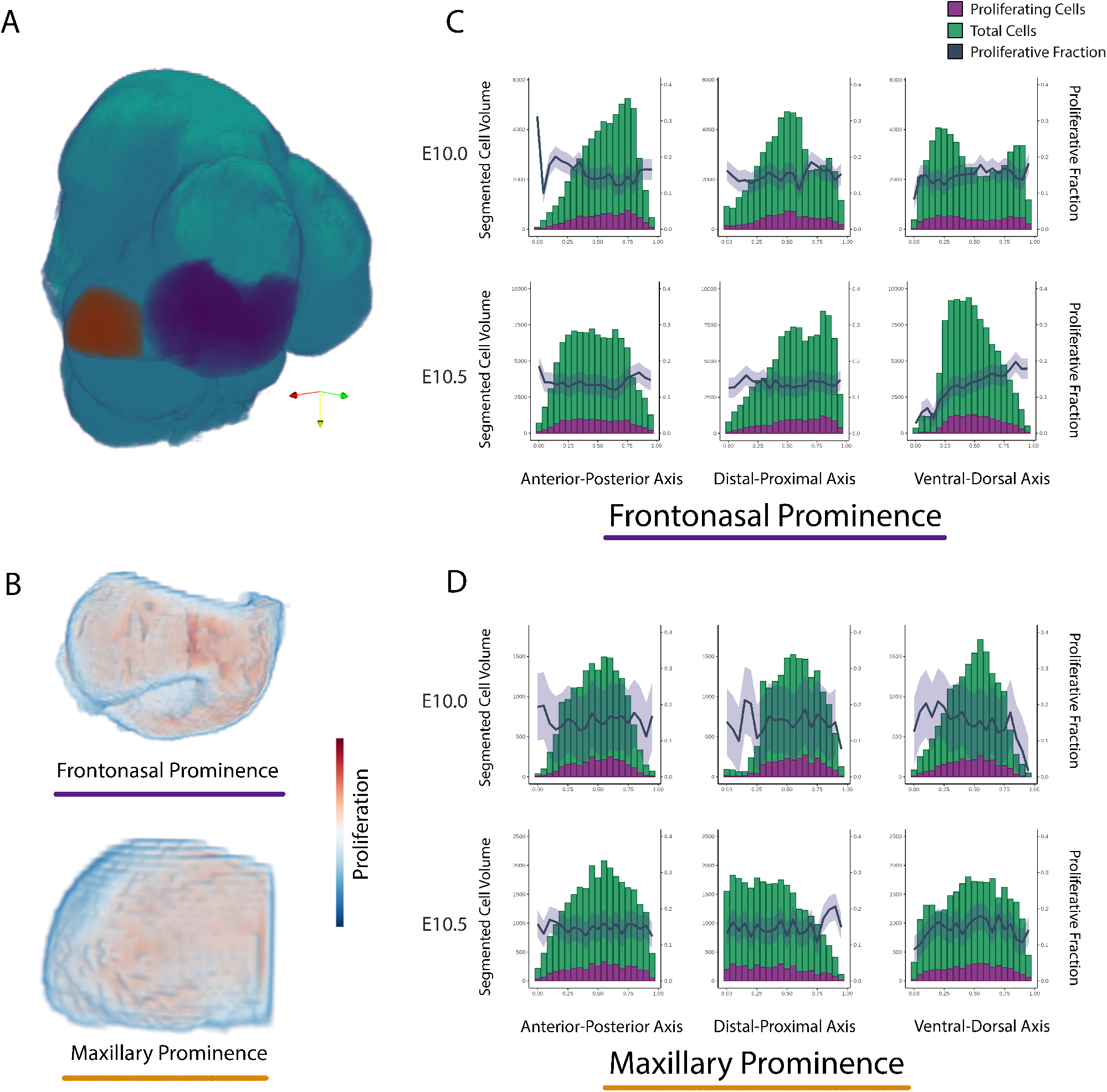
Segmented cell volume and proliferative fraction along each tissue axis at E10.0 and E10.5 in key tissues. (A) Segmentation of the maxillary prominence (purple) and frontonasal prominence (orange) on the average tissue atlas. (B) Proliferative heatmaps within segmented tissues in the same view as (A). (C,D) Cell volumes and proliferative cell volumes are compared along each axis and reveal largely homogeneous proliferation in these averaged volumes. The blue line represents the proliferative fraction, or the proportion of proliferating nuclei to total nuclei, and the ribbon represents the standard error between specimens.

### Correlation of morphology and proliferation

There are many long standing hypotheses predicting that proliferation is a strong driver of morphology during development (Francis-West et al., 1998; Marcucio et al., 2015). To test this hypothesis, it was important to first examine if proliferation and morphological change are correlated. We used voxel-based methods to identify between-group differences and to examine the correlations between shape (morphology) change and proliferation during the phases of rapid growth and morphologica change of the mouse face. Using the atlases shown in Fig. 3, we registered each subsequent age together (eg. E10 to E10.5 and E10.5 to E11) to calculate the facial shape difference (Fig. 5A,B). We then computed a Pearson correlation between the shape difference and the proliferation in the previous age (Fig. 5C). Shape change was largest between the E10.0 and E11 timepoints. The overall correlation rate for the younger ages approached 0.75, implying a strong correlation between shape change and morphological change. By E11.0-E11.5 however, this has decreased to around 0.4. The decrease in the older age is likely due to a combination of decreased shape change between E11 and E11.5 and an overall decrease in proliferation.

**Fig. 5.**
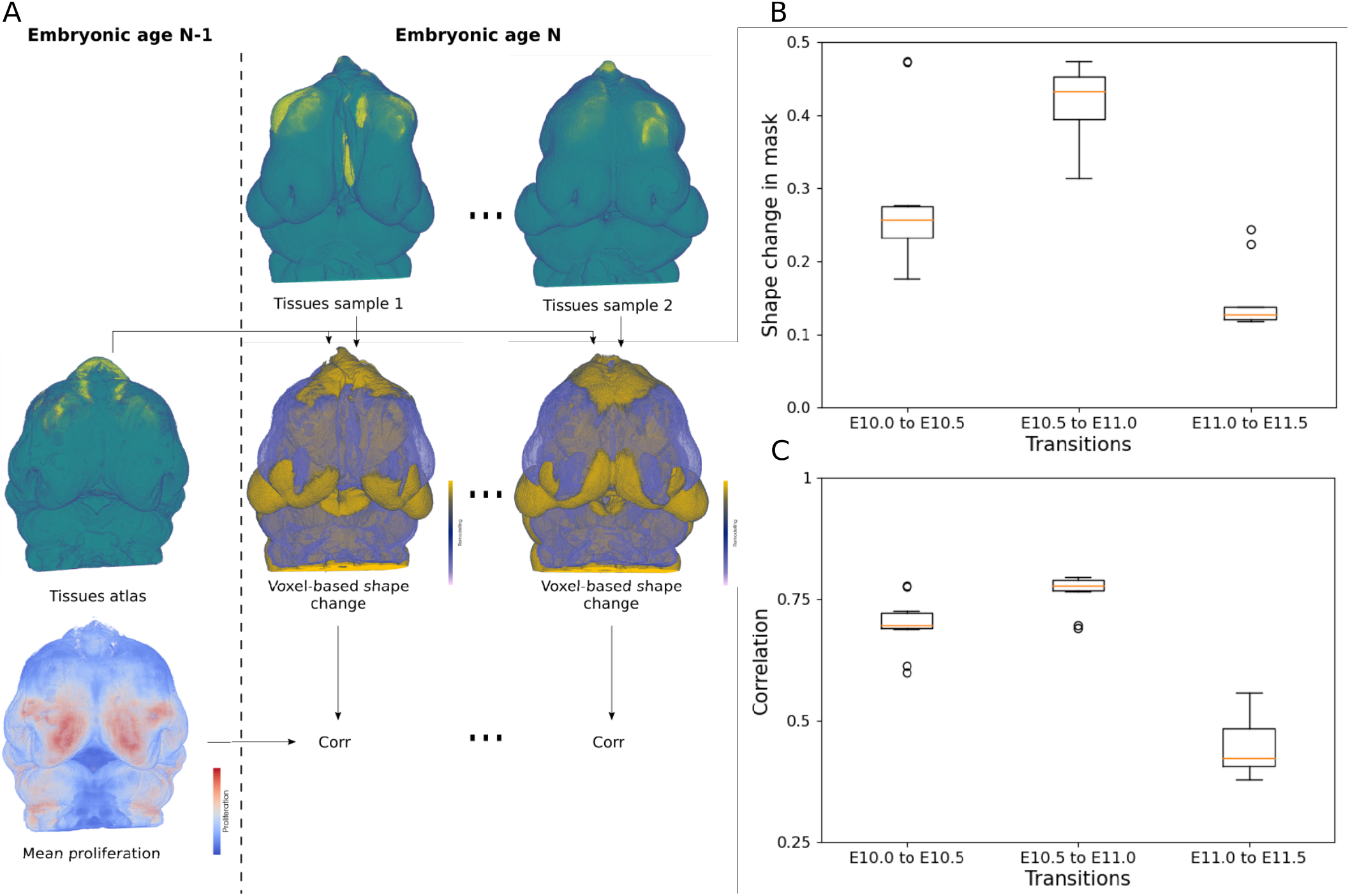
Voxel-based analysis of relation between proliferation and shape change in the face. (A) Workflow to relate the shape and proliferation atlases of the previous embryonic age N-1 to the shape of each of the samples belonging to the present embryonic age N. A differential volume (−1,0,+1) for each sample at age N is obtained by comparing to the mesenchyme of the age N-1 atlas (middle row). Then, each voxel of the differential volume at age N was associated with the corresponding voxel of the proliferation atlas of age N-1. Finally, Pearson’s linear correlation coefficient is computed only in a mask corresponding to the mesenchyme of the face. (B) Box-plots showing the proportion of voxels in the fixed mask of the face that presents outward (+1) or inward (−1) remodeling between the consecutive ages studied in this work (E10.0-E11.5). (C) Box-plots presenting the absolute value of the correlation coefficient between the proliferation at age N-1 with the shape change from N-1 to N, in consecutive ages.

Next, we sought to directly test the hypothesis that proliferation in regions of significant shape change was larger than what could be expected by chance. This hypothesis is important because if proliferation is a driver of morphological change, it would be predicted that regions of increased rates of shape change should also have increased rates of proliferation. In order to test this, we used additional non-linear voxel-based morphometrics methods to identify regions of local tissue expansion or contraction (representing regional morphological changes) between atlases generated from each age group (Fig. 6A). As expected based on the principal component plot, no regions of expansion or contraction were identified as significant between E10 and E10.5 using these methods. We did not detect any stage-related shape effects at E10, likely due to sample size. However, we observed that E11 and E11.5 contained 627,878 voxels (10.4%, p < 0.05, t = 3.23) and 769,245 voxels (12.7%, p < 0.05, t = 3.08) with significant determinants, respectively. The Dice coefficient for the resulting shape significance masks was 0.57, indicating substantial overlap (Fig. 6A). The largest and most consistent shape changes can be traced to the frontonasal prominences, which undergo immense mediolateral expansion towards the midline during mid-gestation. We then compared proliferation in the expanding regions to an equal number of randomly selected voxels (Fig. 6D,E). Interestingly, we observed that the mean proliferation intensity within these shape significance masks fell outside the distribution of randomly sampled intensities for every specimen. While the mean intensity of the randomly sampled proliferation masks at E11 was 57.26, the mean intensity of the shape significance masks was 70.14. This represents a 22.5% increase in average proliferation at sites of major shape change relative to random sites. Similarly, the mean intensity of the randomly sampled masks at E11.5 was 63.23, whereas the mean intensity of the shape significance masks was 84.64. This delta reflects a 33.9% increase in average proliferation. This result shows that there is more proliferation in regions undergoing expansion than could be expected by chance.

**Fig. 6.**
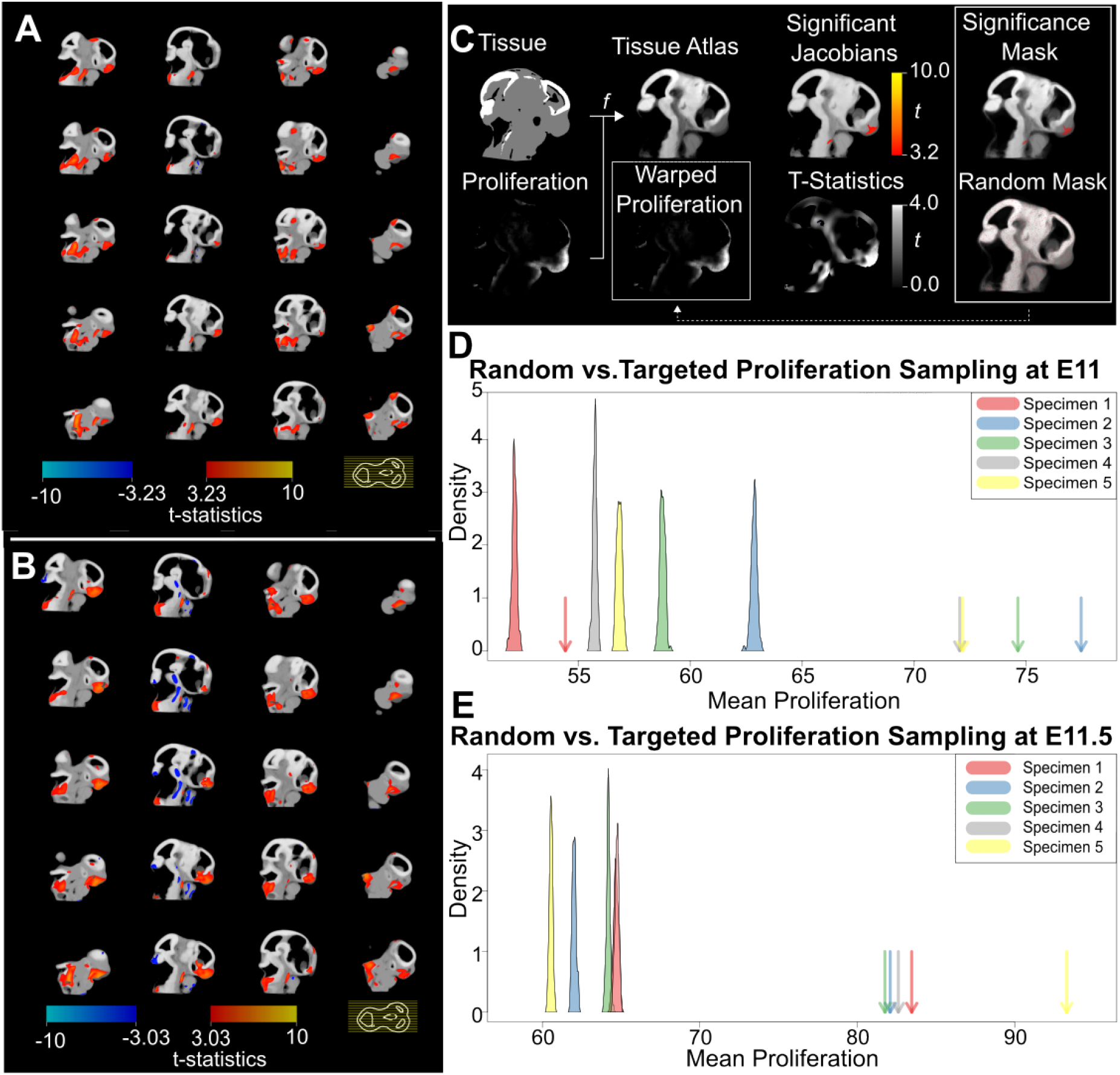
Voxel-based morphometry for whole embryo heads. (A) Statistical parametric map showing sagittal slices of craniofacial regions which undergo significant shape change from E10.5 to E11. B. E11 to E11.5. The slice legend is displayed on the bottom right, with each yellow line corresponding to an individual slice in the volume. Hot colours (red-yellow scale) indicate voxel expansion, whereas cold colours (blue-teal scale) represent voxel shrinkage. (C-E) Localizing the relationship between cell proliferation and shape change in anatomical context. (C) Overview of the image analysis workflow. We non-linearly registered each tissue volume to a novel tissue atlas, and use the corresponding transformation to warp the proliferation volume into the atlas space. Next, we calculated the Jacobian determinants of the tissue registration field and visualized significant stage-related shape deviations (t values) with a parametric map. The red-yellow t value scale indicates levels of increasingly large shape change. Next, we exported a volume of t statistics to generate a shape significance mask (i.e., voxels with a t value above the significance threshold), as well as a set of randomly sampled masks (i.e., voxels with randomly sampled t values). The atlas masks are then overlaid (dotted line) onto each warped proliferation volume to relate proliferation to shape change in the anatomical context. (D-E) Density plots comparing the mean proliferation of N=100 randomly sampled masks (bell curves) against the mean proliferation of the significance mask (arrows) for each specimen. E11 and E11.5 are displayed separately.

To determine whether spatial patterning of proliferation drives ontogenetic variation in morphology, we examined the relationship the 3D volumetric patterning of proliferation and external morphology among individual embryos across the entire age range of the sample. Our initial two block two-block partial least squares regression (PLS) for the entire sample showed a very strong relationship between proliferation and morphology (r = 0.97, z = 2.161, p = 0.001). To determine whether variation in morphology among individuals correlated with the patterning of proliferation, independent of this strong ontogenetic trajectory, we examined the correlation between the residuals for proliferation and morphology on somite stage. This yielded a much weaker and statistically non-significant correlation (r = 0.607, z = 1.507, p = 0.072). However, this analysis is complicated by the clear and strong separation in the patterning of proliferation between the younger and older portions of the ontogenetic range (see Fig. 3B,D). Most of the shape changes associated with proliferation are driven by earlier stages (8-14) and the relationship between proliferation and morphology is different between these two time ranges. Accordingly, we split the sample into the two groups apparent in the PCA plots for both shape and proliferation and then performed PLS regressions within those groups. The resulting 2-block PLS analyses revealed strong relationships between the patterning of proliferation and morphology within each age group (Fig. 7). The relationship was somewhat stronger in younger embryos (8-13 somites) (r = 0.973, z = 3.3.41, p = 0.001; Fig. 7A). Within this sample, shape changes associated with proliferation were mostly localized to the medial nasal region. In contrast, the PLS in the older embryo group (15-20 somites) showed a slightly weaker correlation between proliferation and shape changes (r = 0.91, z = 1.957, p = 0.024; Fig. 7B). Here, shape changes associated with proliferation were mostly associated with maxillary growth.

**Fig. 7.**
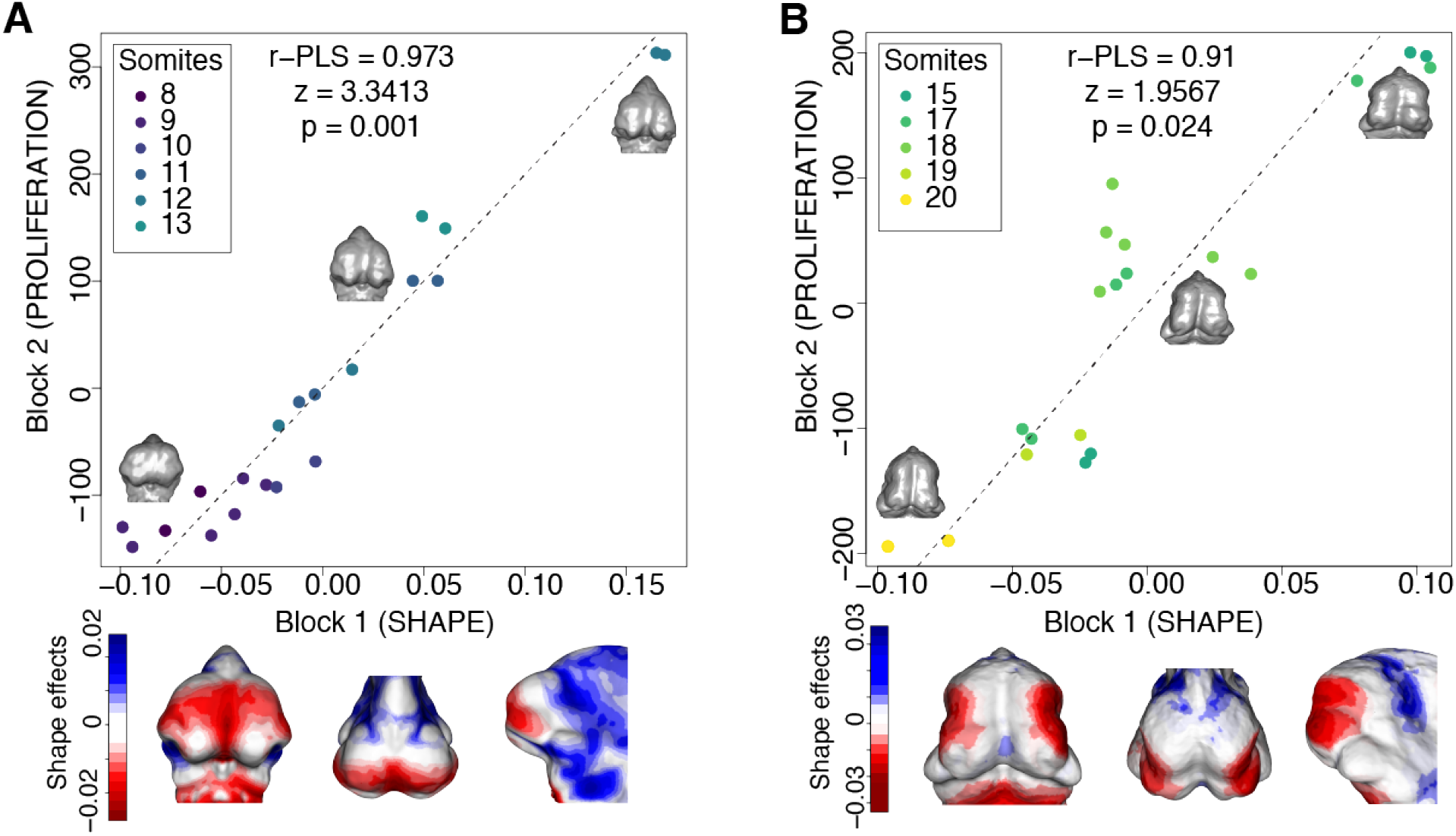
Quantitative relationship between morphology and spatial patterning of proliferation. (A) 2-Block Partial Least Squares (PLS) analysis of Procrustes shape coordinates and proliferation in younger embryos (8-13 somites). Morphs within the scatterplot were constructed from Procrustes coordinates for each specimen. Heatmaps below the scatterplot depict shape changes associated with proliferation, based on the minimum and maximum values along the first latent variable of block 1 in the PLS analysis (shape), which tends toward more medial nasal proliferation. (B) 2-block PLS analysis of Procrustes shape coordinates and proliferation in older embryos (15-20 somites). Morphs were constructed from Procrustes coordinates for each specimen, and heatmaps depict shape changes associated with changes in proliferation along the first axis (latent variable of shape block), which moves more toward the maxilla with increasing age.

## DISCUSSION

Light-sheet microscopy (LSFM) offers new opportunities to investigate cell biology within its 3D context, but these opportunities have been difficult to realize due to challenges of quantifying the outcomes. Here, we present a method to overcome some of these challenges and apply it to quantifying the relationship between proliferation and morphology during the events of primary facial morphogenesis in mouse embryos. We place our results in the context of both understanding the role of proliferation in establishment of facial shape and also of further developing tools to push the field forward.

### Structure of growth

Morphogenetic mechanisms drive not only the large changes in morphology over ontogeny but also the smaller differences in stage-specific morphology among individuals, including structural birth defects. To study the cellular basis for among-individual variation, it is necessary to quantify morphology and cellular dynamics simultaneously in individual embryos. This contrasts to the more common approach of obtaining cellular and phenotypic data in different batches of embryos from the same treatment group or genotype. Excellent methods currently exist for high-resolution quantification of morphology in embryos using computed microtomography (Wong et al., 2015; Hallgrimsson et al., 2015) or optical projection tomography (Sharpe, 2004; 2003). Existing methods also allow quantification of gene or protein expression along with morphology for the same embryo (Xu et al., 2015; Green et al., 2017; Marchini et al., 2021). Quantifying cellular dynamics within such a framework is non-trivial as this requires anatomical localization of cellular parameters and, preferably, accurate segmentation of individual cells in anatomical context.

Altering proliferation is one way, of many, to alter morphology during the development or to generate differences in morphology among individuals at a particular developmental stage (Green, 2022). As organisms grow, the number of cells increases. Deviations from spatially homogeneous proliferation such that some anatomical regions proliferate more rapidly than others, or areas where division tends to occur in a spatially directed manner due to patterning of cell polarity are, in theory, ways to generate shape change during morphogenesis. However, spatio-temporal patterning of cell proliferation is not the only potential cellular mechanism for generating morphological variation in growing embryonic tissues (Green, 2022). Planar cell polarity, for example, can produce morphogenetic changes via directional bias in proliferation (Gray et al., 2011). Boehm et al. (2010) showed using a combination of 3D quantification of proliferation and modeling that such directional cell behaviours are necessary to explain vertebrate limb morphogenesis, whereas our work showed that altering Fgf signalling created dysmorphologies that were associated with disrupted polarity of mesenchymal cells in the face (Li et al., 2013). Cell migration is also critical for morphogenesis and has been studied in many developmental contexts. In the neural crest, for example, multiple mechanisms involving secreted factors, extracellular matrix, and physical forces interact to produce dramatic changes in cell position and gross morphology (Bronner-Fraser, 1993; Shellard and Mayor, 2019). Even here, though, patterning of cell proliferation within migrating streams of neural crest cells plays a role (Kulesa et al., 2008). Finally, physical forces influence morphogenesis at multiple levels ranging from interaction with extracellular matrix (Daley and Yamada, 2013; Linde-Medina et al., 2016), among cells due to regulation of cell adhesion (Lecuit, 2005; LeGoff and Lecuit, 2016), and physical interactions between tissues such as the neural tube and facial prominences during morphogenesis (Marcucio et al., 2015; Parsons et al., 2011). These alternative and potentially interacting mechanisms for morphogenesis suggest the need for development of methods to objectively determine the relative contribution of spatiotemporal variation in cell proliferation to fully understand the cellular bases of morphogenesis.

The morphogenesis of the face is complex, likely more so than the limb, because the prominences must grow in different directions and at rates that allow for them to meet and fuse. When this fails to occur, the result is an orofacial cleft and variable disruption of subsequent facial development (Diewert and Wang, 1992). For these reasons, it seems likely that the cellular processes that underlie this growth must be spatiotemporally regulated within the facial prominences. If this applies to cell proliferation, then it holds that proliferation should not be homogeneously distributed throughout the volume of growing tissue but rather concentrated in those areas that are most rapidly growing. If this is the case, then at each stage of embryonic development as well as across stages, there should be a correlation between the 3D spatial pattern of cell proliferation and morphology. This is the hypothesis that we set out to test with the novel methods presented here.

Our results show clearly that across developmental time (E10 to E11.5) during facial morphogenesis, there is a strong correlation between the spatial pattern of proliferation and morphology. This pattern is not linear across this range, with distinct patterns of shape change and similarly distinct spatial patterns of cell proliferation in the earlier compared to the later part of this range (Fig. 7). When the relationship between shape and the spatial patterning of proliferation is analyzed separately within each of these two time periods, strong relationships emerge.

Given the finding that spatial variation in proliferation correlates with shape across developmental time, it is surprising that we found that proliferation within the growing prominences during the earlier part of this period was not strongly spatially patterned. We had expected, for example, to find elevated proliferation in regions adjacent to known sources of proliferative signals secreted in regions of ectoderm, such as Fgf8 (Crossley and Martin, 1995; Marcucio et al., 2005) and Shh (Hu and Marcucio, 2009). Instead, we observed a fairly homogeneous distribution of proliferation throughout the volume of the facial prominences that one would expect would result in a pattern of isometric increase in volume rather than change in shape over time. There are two potential explanations for this finding. The first is that within a rapidly growing volume, only very small differences in relative rate or proliferation are necessary to produce change in shape. The fact that we observed a correlation between the pattern of proliferation and shape is consistent with this explanation. Alternatively, there may be factors other than proliferation that are influencing facial morphogenesis at this stage such as those discussed above. These explanations are not exclusive and further work that integrates modelling as well as investigation of molecular markers for other aspects of cellular dynamics of growth are necessary to disentangle them.

A role for factors other than proliferation is suggested by our finding that we do not see a statistically significant relationship between spatial patterning of proliferation and facial shape when developmental stage (somite stage) is regressed out. While larger samples might reveal a stronger relationship, our findings do not support the view that spatial patterning of proliferation accounts for differences in morphology among individuals at a given stage. To some degree, this may be an issue of power. If only small deviations from homogeneously distributed proliferation results in morphogenesis, then the differences that account for variation in morphogenesis among individuals would be even smaller. Again, the integration of simulation with these experimental findings is crucial to investigate this possibility.

Another approach would be to perform the analyses shown here on mouse mutants where we suspect that spatial patterning of proliferation is disrupted. Our finding that proliferation is surprisingly homogeneously distributed is relevant here as this means that loss of proliferation from localized regions may still affect morphological development. Studies to probe this hypothesis are ongoing. However, as the relative changes in proliferation are generally small, a relatively larger number of specimens is likely needed to test this hypothesis. The ability to test these hypotheses is to an extent still limited by technology. Imaging at 25x and preferably higher is required to quantitatively evaluate planar cell polarity Alladin et al. (2020); Revenu et al. (2014). While this is technically possible, the resulting images would exceed 1TB per embryo and the registration and processing of those images for whole faces is currently prohibitively resource intensive.

### LSFM as a tool

While LSFM imaging is becoming more prevalent as a tool for developmental biology, there are many challenges that users of this technique must overcome to apply this method more generally. One issue is that beyond a certain size and tissue density, such as mouse embryos older than E8.5, live light-sheet fluorescent microscopy (LSFM) imaging is not feasible as the tissue must be cleared McDole et al. (2018). Clearing and sample preparation presents significant challenges, although there is an increasing diversity of clearing techniques available. LSFM imaging has even been used to image full juvenile mice after clearing Tian et al. (2021). Data processing is also a significant challenge as computational time increases exponentially with resolution or object size at the same resolution. One crucial processing step is the accurate identification of individual cells within an image in order to quantify a cellular marker in relation to the total number of cells. Our team has had recent success retraining a convolutional neural network (CNN, U-net) to process embryo LSFM images Lo Vercio et al. (2022). We developed a method that efficiently segments tissues, nuclei, and proliferating cells in lightsheet images of whole embryonic heads from E9.5-E10.5 mouse embryos while also dealing with minor LSFM artifacts, such as blurriness or tissue depth variation. Nucleus-level analysis becomes more challenging as the size of the tissue sample or embryo imaged increases as image artifacts become more pronounced and complex with increasing size.

In this work, the large size of the whole E10-E11.5 embryo head required performing the detailed analysis of proliferation rates within the facial prominences and the whole head (Fig. 4). These regions have low probability of segmentation errors and the manual or semi-automatic isolation of these exterior structures can be readily assessed and corrected by a human observer. Segmentation errors happen for two main reasons. The first is that staining and clearing tends to work better close to the exterior of the sample, which results in depth of tissue-related artifacts. Similarly, whole-mount immunohistochemistry is affected by tissue depth due to variation in antibody penetration Yokomizo et al. (2012) and this is apparent as loss of signal in interior structures. For this reason, total nuclei counts are included only in the analysis of the facial prominences for the earlier part of the studied age range (E10-E10.5), while total proliferation data was used more extensively in this work. Further, as we approach the resolution limits of individual nucleus detection, apparent merging of nuclei will bias the segmentation process. For this reason, we used a voxel-based approach rather than a single-cell approach for the quantification of the proliferative fraction (Fig. 4). Also, each proliferation map was normalized before computing the proliferation atlases (Fig. 3A) and performing individual-level analyses (Fig. 3D and Fig. 7). This was to reduce potential effects of inter-sample variability of the proliferation marker. Further work in improving staining, clearing, and nuclei and proliferation segmentation is needed to increase the detail and accuracy of the 3D models constructed within the proposed framework.

To quantify cellular dynamics in the morphological context, it is necessary to register each volumetric image to a common atlas in order to place both morphology and the spatial distribution of cells into a common shape space. Atlas generation for early mouse development is a computational challenge due to the high amount of shape variation among embryos Wong et al. (2015); Devine et al. (2020; 2022). Improvements to atlas generation will require larger sample sizes as well as tighter control of developmental time by increasing the representation per somite stage or by more refined methods of staging. In this work, the more pronounced image artifacts (blurriness, signal loss) in the older embryos (E10.0-E11.5) resulted in poorer performance for the U-nets for tissue segmentation compared to E9.5-E10.5 embryos. This occurred even when U-nets were re-trained with images from our data. As a result, user intervention was required to correct 3D tissue models in cases where tissues were misclassified. Current work is focused on solving or mitigating these issues so that we can apply non-linear registration methods that have less tendency to introduce noise during registration to the atlas Devine et al. (2020). This will lead to more accurate atlases, better quantification of proliferation maps, and improved ability to handle large and shape-diverse samples of embryos.

### Conclusion and Future Directions

This work represents a substantive step in the direction of building the toolkit necessary for quantitative integration of analyses across multiple levels of genotype-phenotype maps. While there are existing methods to quantify gene-expression, cellular dynamics, histology, and external morphology in isolation, there are relatively few methods that enable the simultaneous quantification of more than one of these levels in individual embryos. Such efforts are critical in order to create a mechanistic understanding of the generation of phenotypic variation by developmental processes. While many challenges remain, the method presented here can support the systematic analysis of cell proliferation and morphology in complex morphogenetic contexts such as the development of the vertebrate face. Future efforts will refine this method but also find ways to incorporate additional levels such as gene expression, spatial transcriptomics, and other multiple-molecular marker methods within similar anatomical registration-based frameworks.

## MATERIALS AND METHODS

### Clearing and staining

Dams were sacrificed by isoflurane overdose, and embryos were harvested between E9.5 and E11.5. Following harvest, embryos were washed in PBS for 30 minutes and fixed overnight in 4% paraformaldehyde (PFA) at 4°C. Embryos were cleared with a modified CUBIC protocol to remove lipids and heme while preserving morphology, based on Susaki et al. (2015). Only typically developing embryos were used for further examination.

Following overnight fixation, samples were washed for 24 hours. in PBS to remove blood. Samples were serially dehydrated in increasing concentrations of methanol in PBS (25%, 50%, 75%, 100%) for 30 minutes each. Samples can be stored long-term at this point in 100% methanol at -20°C. Samples were permeablized in 5% H_2_O_2_/ methanol overnight at 4°C. Samples were rehydrated in decreasing concentrations of methanol in PBS (100%, 75%, 50%, 25%) for 30 minutes each, followed by a 2 hour wash in PBS. Samples were incubated overnight in 1:1 PBS:CUBIC-1 (25% urea, 25% Quadrol, 15% Triton X-100), then in 100% CUBIC-1 until transparent (approx. 3-5 days), shaking at 60 rpm at 37°C. Once transparent, samples were rinsed twice in PBS for 2 hours, then once more overnight to stop clearing. Samples were then blocked for 36 hours in 5% Goat Serum, 5%DMSO in 0.1% Sodium Azide/PBS at 37°C to prevent nonspecific binding.

For immunostaining, samples were incubated in 1:200 conjugated antibody (Anti-Histone H3 (phospho 10) antibody (Alexa Fluor 647)) for proliferation staining, 1:4000 Nuclear Green for nuclear staining, and 5% DMSO in 0.1% Sodium Azide/PBS for 5 days at 37°C. Following staining, samples were washed several times in PBS-Tween 0.5% to remove excess antibody. Samples were beheaded, and heads were embedded in 1.5% low melting point agarose and incubated in CUBIC-2 (25% urea, 50]% sucrose, 10% triethanolamine) for 24 hr. Tails were also embedded to stage embryos after imaging.

### Imaging

Light-sheet images of agarose-embedded samples were obtained with a Zeiss Lightsheet Z.1 microscope. Heads were imaged at 5 *μ*m intervals with 0.9 *μ*m resolution with a 5X objective in CUBIC-2. Tails were separately imaged at 7 *μ*m intervals with 2.5 *μ*m resolution for manual counting of tail somites as a measure of embryonic age (Chan et al., 2004). Image stacks were taken with a 10% overlap and stitched together in ZEN Blue (Zeiss).

### Automatic analysis workflow

#### Segmentation

The U-net for tissue segmentation was trained using Nuclear Green images (1024 *×* 1024 downsized to 256 *×* 256), 21 images from E11.0 embryos and 10 images from E11.5 embryos. For a test set of 8 images from a E11.0 embryo and 10 from a E11.5 embryo, the U-net reached an accuracy of 0.806 and a Dice score of 0.727. The mesenchyme and neural ectoderm of the 20 E10.0-E11.5 embryos analyzed in this work were segmented using this U-net, while the nuclei and proliferating cell segmentation was done using the U-nets and ImageJ-Fiji plug-in described in Lo Vercio et al. (2022). The segmentation volumes were transformed to isotropic dimensions via shrinking of the XY dimension, and corrections for the segmentation volumes of tissues (mesenchyme and neural) were performed in 3D Slicer (Kikinis et al., 2014) by a human observer, particularly removing deposits outside the sample not segmented as background, and correcting voxels misclassified as neural ectoderm to the mesenchyme class.

#### Registration for atlases

To build general shape and proliferation maps at each developmental age (E10.0, E10.5, E11.0 and E11.5), one sample per age was selected as reference. Then, it was manually oriented, and anything below the mandible was deleted from the volume. The five samples of each stage were mirrored to double the amount of data. Then, a landmark-based rigid registration was performed between each of the remaining samples in the age group and their reference sample, using five landmarks placed in the head by an expert observer (Devine et al., 2022) via SimpleITK in Python (Lowekamp et al., 2013)). Then, the resulting transformation was applied to the correspondent tissue segmentation. A groupwise-affine registration was performed among the ten registered tissue segmentations using SimpleElastix in Python (multiresolution registration with 5 number of resolutions, 30000 maximum iterations, linear interpolator, nearest neighbor interpolator for resampling) (Marstal et al., 2016; Devine et al., 2020). Finally, the voxels of the shape atlas were labelled as background, mesenchyme, or neural ectoderm using majority voting, implemented in MATLAB (Fig. 3A).

The chain of rigid and affine registrations of each sample was latter applied to the corresponding volumes of nuclei and proliferating cells. The mean proliferation per voxel was computed using a 0.15mm*×*0.15mm*×*0.15mm window. This proliferation map of the sample is normalized using percentile-based equalization Weigert et al. (2018). The mean proliferation map for each age was obtained by averaging the proliferation maps of the five embryos and their mirrored maps (Fig. 3A).

#### Registration for bulk analysis

To analyze the shape and proliferation variation among all the samples used in this work, 37 landmarks were placed in the face of the 20 samples and their mirrored volumes by an expert observer (Percival et al., 2014). Then, a landmark-based similarity registration (translation, rotation, and scaling) was performed using VTK in Python (Schroeder et al., 2006) between each sample and one E10.5 sample used as the reference (transformed as in the previous section). These transformations were latter applied to the corresponding proliferation maps.

#### Voxel-based study

Using the tissue and proliferation maps produced by the latter described registration process, we proceeded to extract the shape and proliferation in the face of the registered embryos. To build this mask, the forty mesenchyme segmentations were extracted and voxels labelled as mesenchyme more than 30% of the time were assigned to a primary mask. Then, voxels further away than 0.39 mm from the surface of this mask were excluded. Finally, an expert observer manually refined the mask using 3D Slicer leaving only voxels belonging to the face. With this mask, the proliferation map values were extracted for the forty samples.

#### Landmark-based study

We analyzed and visualized facial shape using geometric morphometrics methods (GMM). We performed a Generalized Procrustes Superimposition Analysis (GPA; Gower (1975)) to extract the aligned Procrustes shape coordinates from the manual landmark dataset (to focus on shape differences only on the face and not the whole head; Fig. 3C), using the package *geomorph* (Adams et al., 2022) in R (R Core Team, 2020). The Procrustes shape coordinates represent each specimen’s shape, and we ordinated these with a Principal Components Analysis (PCA), to visualize the axes of maximum shape variation and its association throughout ontogeny (using tail somite number as a proxy for age), with the R package *geomorph* (Fig. 3B). We also performed a PCA on the flattened matrix from the proliferation array, to be able to visualize differences in proliferation associated with age (Fig. 3D).

To assess the degree of association between face shape and proliferation patterns, we performed a two-block Partial Least Squares (PLS; Rohlf and Corti (2000)) analysis using the R packages *Morpho* (Schlager, 2017)) and *geomorph* (Adams et al., 2022). PLS latent variables were calculated as the linear combinations of the Procrustes shape coordinates (Block 1) and the flattened proliferation matrix (Block 2), which maximized the covariance between the two blocks. We plotted an ordination PLS scatterplot, coloured by the number of tail somites (indicating age), displaying the first latent variable for both the shape dataset (Block 1) and the proliferation dataset (Block 2), using *geomorph* (Fig. 7). We split the embryonic dataset into two age-relevant groups, based on their distinct grouping in both the facial shape morphospace (Fig. 3C) and proliferation patterns (Fig. 3D). Meshes to produce morphs were obtained from smoothed atlases using morphological closing. Finally, we generated morphs using the shape changes associated with proliferation changes, using the R package *Morpho* (Fig. 7).

#### Single tissue analysis

Based on our whole-head proliferation atlas, the frontonasal and maxillary prominences were identified as the most actively proliferating mesenchymal tissues in the face between E10.0 and E11.0, and because of that were selected for tissue-specific analysis. Tissues were segmented on the atlas in 3D Slicer and back-propagated to each sample in the group. Total and proliferating nucleus volumes were calculated by segmenting cells as described above using a connected components method. Nuclear volumes were used instead of counts to reduce segmentation errors in dense tissues, especially in the Nuclear Green stain. Neighbouring cells in dense stains tend to be under-segmented; by summing total volumes of segmented nuclear voxels, there is no need to identify which cell they belong to, removing a potential source of error. Cells below the 1st and above the 99th percentile in volume were considered segmentation artifacts and excluded from analysis. Cells were binned by relative position along each tissue axis, and proliferative fraction was calculated as the proportion of cell voxels that also contained a proliferation voxel in each bin. To identify nonuniform trends in proliferation, a Kolmogorov-Smirnov test of uniformity was conducted on each axis.

## Acknowledgements

The authors would like to thank Nicholas Hanne and Nathan Young (UCSF), James Cheverud and Fernando Andrade Oliveira (Loyola University Chicago).

## Competing interests

The authors declare no competing or financial interests.

## Contribution

Conceptualization: R.M.G., L.D.L., J.C.B, W.D, B.H., R.S.M.: Methodology: R.M.G., L.D.L., B.H., R.S.M., N.D.F., J.C.B; Analysis: L.D.L., A.D., J.D., S.R., M.V-G, R.M.G; Sample preparation: R.M.G., E.B., M.M., M.V-G, A.D., S.G., X.Z., M.B.S.; Software: L.D.L., S.R., A.M., M.V-G; Writing and Revision: R.M.G., L.D.L., B.H., A.D., M.V-G; J.D, J.C.B, W.D., A.L. Visualization: L.D.L., A.D., J.D., M.V-G; Supervision: B.H., R.S.M., N.D.F.; Funding acquisition: B.H., R.S.M., A.L., N.D.F.

## Funding

The work was supported by NIH NIDCR R01-DE019638 to R. M. and B. H., NSERC Discovery Grant ##238992 to B. H., a CIHR Foundation grant to B. H. and R. M. a CFI grant #36262 to BH as well as the Alberta Childrens Hospital Foundation. L.L. was supported by an Eyes High postdoctoral fellowship (University of Calgary), R.G. by a CIHR fellowship, M.V-G by an Alberta Children’s Hospital Research Institute Postdoctoral Fellowship and Alberta Innovates Postdoctoral Fellowship in Health Innovation, M.M. by a Cumming School of Medicine and McCaig Institute postdoctoral fellowship, E.B. by a Cumming School of Medicine and McCaig Institute postdoctoral fellowship, A.D. by a CIHR CGS-M studentship, A.M by an NSERC USRA, M.S. by an NSERC USRA, and N.F. by the Canada Research Chairs Program. This research was enabled in part by support provided by WestGrid and Compute Canada (www.computecanada.ca).

## Data availability

Insert the Data availability text here.

## Supplementary

Insert the supplementary text text here.

## Notes

### Competing Interest Statement

The authors have declared no competing interest.

